# West Asian sources of the Eurasian component in Ethiopians: a reassessment

**DOI:** 10.1101/694299

**Authors:** Ludovica Molinaro, Francesco Montinaro, Toomas Kivisild, Luca Pagani

## Abstract

Previous genome-scale studies of populations living today in Ethiopia have found evidence of recent gene flow from an Eurasian source, dating to the last 3,000 years^1,2,3,4^. Haplotype^1^ and genotype data based analyses of modern^2,4^ and ancient data (aDNA)^3,5^ have considered Sardinia-like proxy^2^, broadly Levantine^1,4^ or Neolithic Levantine^3^ populations as a range of possible sources for this gene flow. Given the ancient nature of this gene flow and the extent of population movements and replacements that affected West Asia in the last 3000 years, aDNA evidence would seem as the best proxy for determining the putative population source. We demonstrate, however, that the deeply divergent, autochthonous African component which accounts for ∼50% of most contemporary Ethiopian genomes, affects the overall allele frequency spectrum to an extent that makes it hard to control for it and, at once, to discern between subtly different, yet important, Eurasian sources (such as Anatolian or Levant Neolithic ones). Here we re-assess pattern of allele sharing between the Eurasian component of Ethiopians (here called “NAF” for Non African) and ancient and modern proxies area after having extracted NAF from Ethiopians through ancestry deconvolution, and unveil a genomic signature compatible with population movements that affected the Mediterranean area and the Levant after the fall of the Minoan civilization.

## Results and Discussion

To determine the most likely source of the Eurasian gene flow into the ancestral gene pool of present-day Ethiopians we have used a combination of ancestry deconvolution (AD) and allele sharing methods^6^. AD refers to analyses that determine the likeliest ancestry composition of genomes of individuals with mixed ancestry at fine haplotype resolution. These methods have allowed us to i) exploit high quality modern data and ii) harness the power of allele sharing tools on genetic fractions with no or reduced African contributions. Such a strategy, while potentially beneficial, introduce a novel source of bias which we aimed to explore here. Particularly, after AD of 120 Ethiopian genomes^7^, we assigned each genomic SNP into one of the following four categories based on the method likelihoods (see Methods for further details): 1) confidently non African (NAF); 2) low confidence non African (X); 3) low confidence African (Y) and 4) confidently African (AF, consistently filtered out from our analyses). While basing our inference on the NAF component alone, we here demonstrate that the component X does account for a minority of the genome and, when analysed together with NAF does not qualitatively change the results. Furthermore, when joining together the NAF and AF confidently assigned components (to create “Joint” components) we recapitulate the signals of the global population (prior to ancestry deconvolution), showing that the X and Y components are not holding a considerable or peculiar genetic signature and hence ruling out, in this study, the role of ancestry deconvolution as a potential source of artifacts. For the sake of clarity, out of the four admixed Ethiopian populations available from Pagani et al. 2015 (Amhara, Oromo, Somali, Wolayta), we report results only on the NAF component of Amhara. Comparable results for the other three populations, which we chose not to lump into a heterogeneous Ethiopian super-population to emphasize potential population-specific peculiarities, are provided in Supplementary Information.

A preliminary exploration of the NAF genomes through ADMIXTURE (Figure S5) and projected PCA showed them to fall within the range of Eurasian populations, close to ancient populations with a high Anatolian Neolithic component (e.g. Anatolia_N and Minoans) (Figure 1 and S1-S4). Notably, several Jewish populations from North Africa cluster with NAF as well. The affinity between Anatolian Neolithic and NAF was further highlighted by *f3* out-group statistic, in contrast to results obtained with the genomes before ancestry deconvolution (Supplementary Figure S6). Overall, whole-genome sequences of all the Ethiopian populations appear closer to ancient Near Eastern populations such as: Minoans, Natufian, Levant Neolithic and Anatolian Neolithic. On the other hand, their NAF components appear closer to populations with a high Anatolian rather than Levantine (such as Minoans, Sardinians and Anatolia Neolithic) component. The highest genetic affinity to the NAF components was observed among North African (Tunisian, Libyan and Moroccan) Jews (See Figure S6), as already seen in the PCA clustering (See Figures 1, S1-S4).

**Figure 1:**
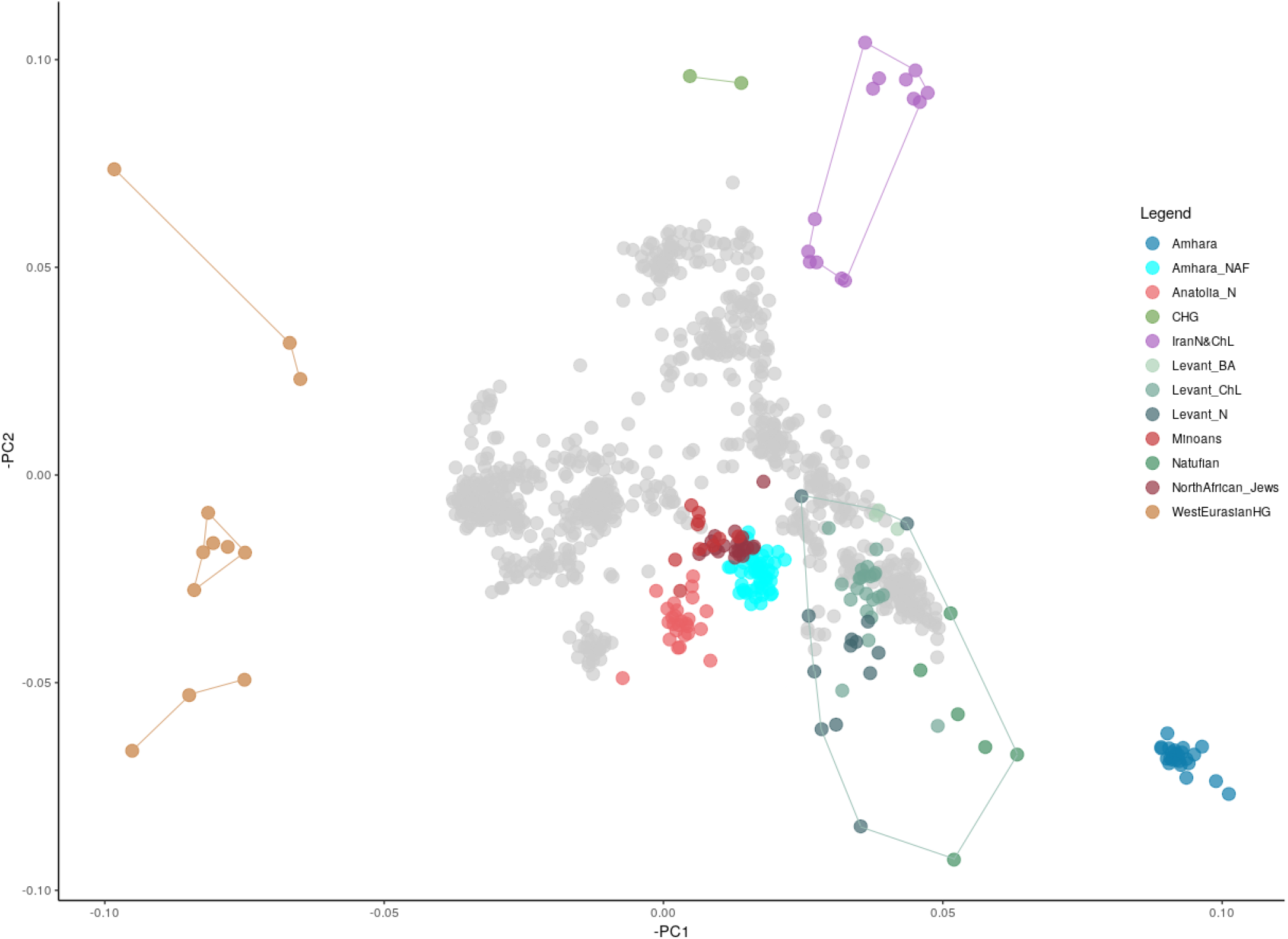
Principal component analysis of modern West Eurasian populations used as a scaffold (grey points) on which we projected ancient and ancestry deconvoluted genomes. To highlight the populations studied we coloured European hunther-gatheres, ancient genomes from Anatolia and Levant areas, Jews from North Africa and Amhara whole and NAF genomes. Variance explained by PC1 is 0.9% and PC2 is 0.3%

We further dissected the observed affinity between NAF and Anatolian Neolithic-like populations through a set of *f4* tests aimed at refining through more and more stringent comparisons the best proxy population for the Eurasian layer (Figure 2). The whole-genomes, with both African and Non-African component, are significantly closer to a Levantine ancestry rather than Anatolian (Z-Score 2.98), with them being closer to Levant_ChL individuals than Levant_N. On the other hand, NAF is shown to be closer to a Neolithic ancestry from Anatolia rather than any Levantine one (Z-score −2.847) and, among Levantine populations, notably closer to Levantine Chalcholitic than to Bronze Age groups or contemporary Lebanese. We further compare the best proxies for the Non African component using the top scoring populations from Outgroup *f3* analyses. Minoans appear to be as close to NAF as Anatolian Neolithic individuals (Z-Scores < 1). When we delved into the North African Jews signals, they broadly show affinity with NAF with particular reference to Jews from Tunisian. Similar trends were observed for all other Ethiopian populations (Figure S7 and Table S1) and did not change when considering alternative combinations of deconvoluted components (Figure 2). Given that our ability to pinpoint the actual source of the NAF component is inherently limited by the availability of ancient and modern populations, we used qpGraph (Supplementary Figures S8,S9 and S10) and qpAdm to model NAF as a mixture of the major axes of genetic diversity that best described the Mediterranean area at the time of the studied event, following Lazaridis et al. 2016. When looking at the global genomes, our qpAdm results replicate a Levant_N origin for the Eurasian component of Ethiopians^3^ (Figure 3, left column). For further results on the other Ethiopian populations see Table S2 and Supplementary Figure S11. In sum, similarly to Minoan and Tunisian Jewish populations, the non African component of Ethiopian populations can be best modelled as a mixture of ∼85% Anatolian_N and ∼15% CHG composition of ancestries (Figure 3, columns 2,3,4).

**Figure 2:**
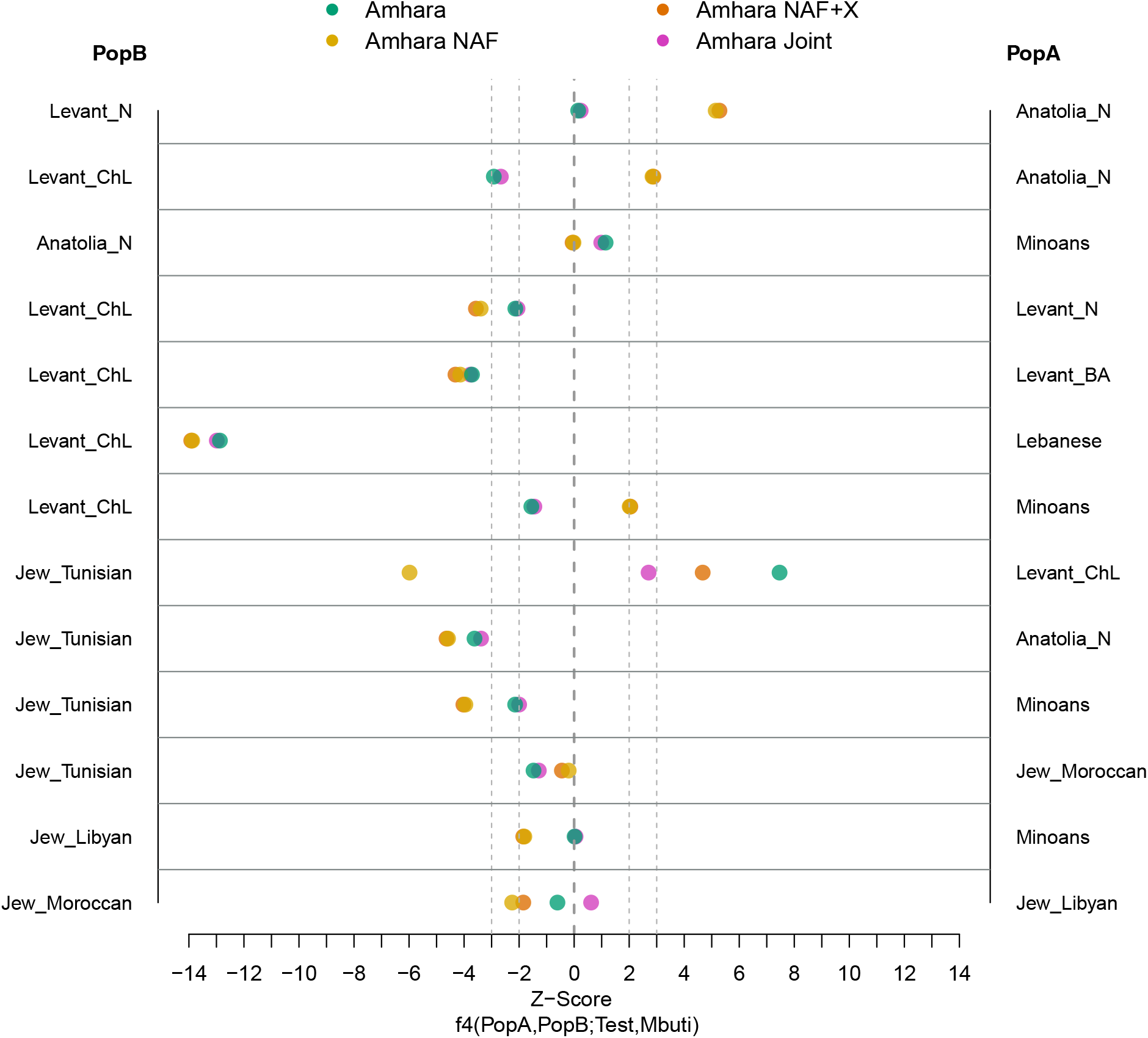
*f4* statistic test on Amhara in form of (PopA,PopB;Test,Mbuti) to assess genetic similarity between Amhara and respective NAF genomes to pairs of several Near Eastern populations. A and B populations are listed in the left and right side of the plot, respectively. Values in x axis indicate the Z-Scores, we draw two lines to highlight |z-Scores|= 2 and 3. Points with |z-Score| > 3 indicate a clear affinity of the test population towards one of the other population. Amhara’s segments tested: Amhara whole-genome (Amhara, in blue), the Non African component (Amhara NAF, in yellow), Amhara African and Non African components together (Amhara Joint, in violet) and Amhara NAF with X component (Amhara NAF+X, in orange).

**Figure 3:**
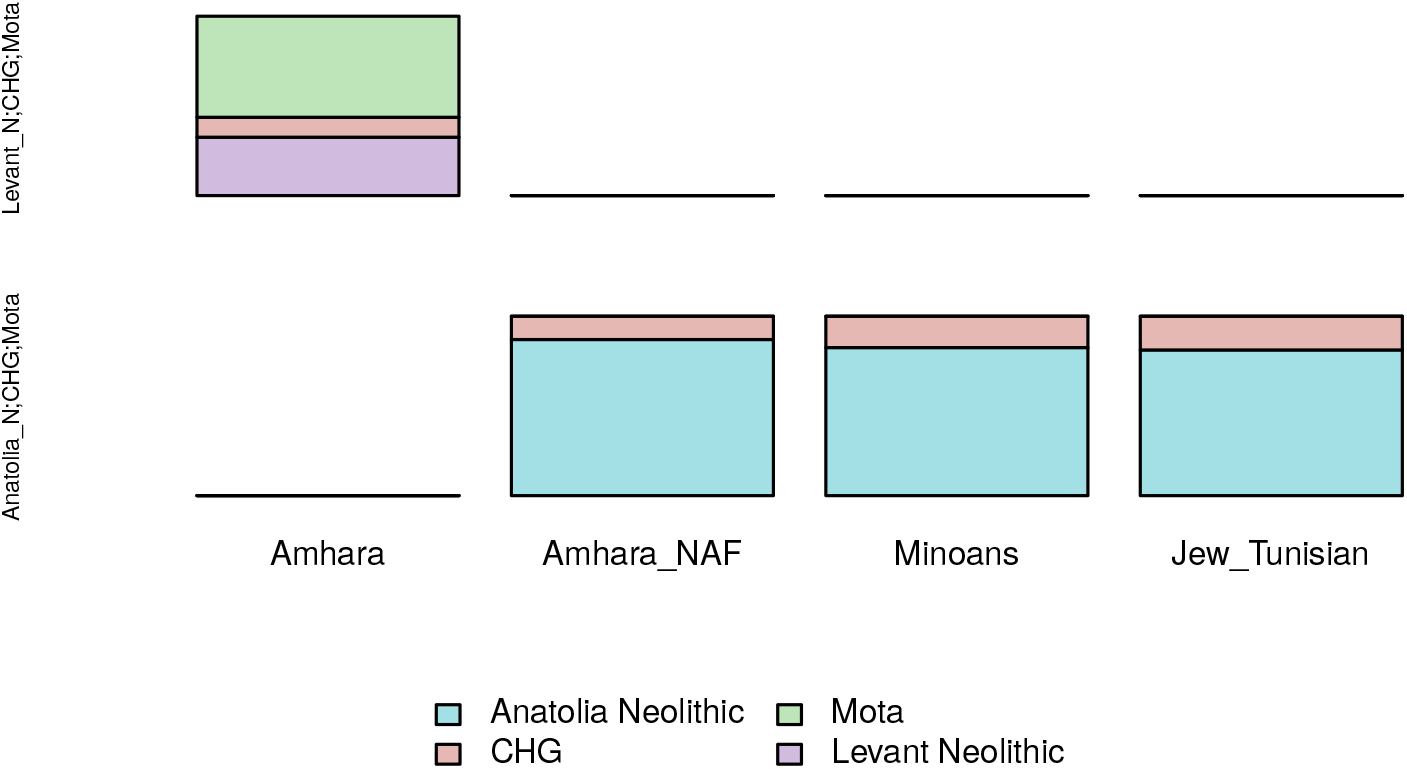
Modelling Amhara, Amhara_NAF, Minoans and Jews from Tunisia as a mix of Mota and Near Eastern populations, with 2 and 3 ways admixtures. Violet indicates the Levantine component, pink the Caucasus Hunter-Gatherers, light green the African component and light blue highlights the Anatolian ancestry. The left side of the graph lists the sources used to model the populations in the x axis; unfilled boxes indicate unfeasible results or p-value < 0.01.

While this mixed ancestry component likely reached Ethiopia only within the last 3,000 years, these results should not be interpreted as involving a direct connection or descent line between Neolithic Anatolia and Ethiopia. Instead, these results can potentially be seen as informative for the identification of candidates among the available ancient and modern populations which, following geographic and chronological considerations, may be suitable proxies for one or more populations that mediated the Eurasian gene flow to East Africa. Of the ones analyzed here, Minoans and Tunisian Jews seem to provide the two closest matches to NAF, adding on top of the genetic evidence a criteria of space/time compatibility. A tentative links between these three groups may be provided by the maritime trade routes connecting Crete (home to the Minoan culture) to the Levant^8,9,10^ and by the shuffling role played by a horde of nomads who navigated throughout the Mediterranean Sea 3 kya: the Sea People. These tribes left traces of their passage both in Crete, in Anatolia, when they fought the Hittite Empire and in Egypt and the Levant, and are told to have settled in the land of Canaan, known also as Palestine^11^. Interestingly, among those tribes that settled in Palestine there were: Denyen, Tjeker and Peleset. Although there are different theories around the origin of each of the tribes, there are suggestions that link the Denyen with the tribe of Dan, from which Jews from Ethiopia have been said to descend and Peleset to their neighboring Philistines^12^. The role of Sea People may therefore be crucial in explaining a temporary presence of a Minoan-like ancestry in the Levant, bringing Anatolian-like components to levels as high as 85%. A pulse of populations with Anatolian-rich ancestry has just been recently detected in Iron Age Levant, appearing and disappearing from the archaeological record within a range of few centuries^13^. Our results offer a solution to this disappearance, given that their signal may have become erased as a consequence of major warfare after 1000 BCE^14^, bringing these genetic components towards Ethiopia and North Africa.

In conclusion, our work shows that when the mixing components are deeply differentiated, such as in the case of contemporary Ethiopians, ancestry deconvolution increases the sensitivity of allele sharing tests and enables to fully exploit the high quality of modern genomes.

## Acknowledgments

The authors would like to thank Dr. Doron Behar and Dr. Iosif Lazaridis for fruitful discussion on an early version of this manuscript. This work was supported by the European Union through the European Regional Development Fund Project No. 2014-2020.4.01.16-0024, MOBTT53 (LM, LP).

## STAR Methods

### Dataset and Samples

We merged different datasets available containing both ancient and modern DNA, African and Eurasian populations from the following publications^15,16,17,3,18,19,20,21,5,22,23,24,25,26,27,28^. Northeast African populations whole-genome sequences were taken from Pagani 2015^7^, and included 5 modern Ethiopian populations: Amhara, Gumuz, Oromo, Somali and Wolayta. We chose to focus on the whole genome sequence data rather than on SNP arrays^1^ to increase the number of available SNPs to be compared with aDNA and other references. To maximize the number of individuals typed at each SNP, we downsampled the dataset to 1037084 markers to match the ones of Human Origin Array on which most of the ancient DNA samples were typed. For ease of exposition we chose Amhara, the population with the highest Eurasian fraction among the available ones^7^, to represent all main results. We provide full description of all other Ethiopian populations in Supplementary Material. Similarly, we chose not to group all the available samples within a single “Ethiopian” population, to allow for group-specific stories to emerge.

### Ancestry Deconvolution

#### Subsetting Modern Genomes

From phased genomes, we refined the ancestral components identification in Eastern Africans individuals provided by Pagani 2015 with PCAdmix^29^. For every 20 SNPs window of the genome, there is a probability for the window to have a source of African (AF) ancestry or Non African (NAF) ancestry (in which case the probability is 1 – AF), which is given by fbk values and refined with Viterbi algorithm^30^. We set a fbk threshold of 0.9 probability in order to assign every window to either one layer of ancestry or the other. If a window did not reach the threshold for any component, it would have been labeled as unassigned. CEU (Utah residents with ancestry from northern and western Europe) were used as a proxy for the Non African component, and Gumuz (the Ethiopian population showing minimal introgression) were used as a proxy for the African component following Pagani et al. 2015. Once the ancestral components were detected, we created the "Genomes Subsets" using the windows that reached the threshold. The "Genomes Subsets" are genomes in which for every haplotype only the confidently assigned African or Non African component is retained, while the rest is assigned as “missing data”. Therefore, they are partial genomes in which only the sequences derived from a specific ancestry (either African or Non African) are present (see Yelmen et al. 2019 for further details). The ancestry deconvolution process has been applied to East African populations only from Pagani 2015 populations, namely: Amhara, Gumuz, Oromo, Ethiopian Somali and Wolayta.

#### Sifting through all possible ancestry fractions

To test for possible biases introduced by using CEU as proxy for the Non African component, we further divided the deconvolution results into different segments to investigate specifically the parts of the genome that were not assigned to either ancestry. We retrieved the different components from the fbk values alone, without refining them with the Viterbi algorithm, to maintain all possible segments information. For each of the two ancestries we obtained two components: X and Y, which held the sequences assigned with 51-90% and 10-50% respectively, representing the unassigned sequences in the masking process. The component X is made of sequences that were not assigned to NAF, representing the unassigned segments that we expect to bear Eurasian traces along with spurious African ones; the component Y is made of segments which we expect to be characterized mainly by African traces. The X and Y segments correspond each for 7% of the genome, and we expect their contribution to the final the results to be minimal.

### Principal Component and ADMIXTURE Analyses

We performed PCA as an initial screening method on the dataset with smartpca from EIGENSOFT^31,32^, using the lsqproject option and autoshrink:YES. We used modern European and Near Eastern populations with minimal missingness (–geno 0.1 with PLINK^33^) to compute PCs and projected the rest of the samples included the ancient samples and the Ethiopian NAF genomes. We used ADMIXTURE^34^ software to perform supervised clustering of ancient and decolvoluted genomes using as reference modern European and Near Eastern genomes along with Yoruba as African, Gumuz as East African and Han as East Asian. We used R and ggplot2 package for visualization^35,36^.

### Frequency-Based Allele-Sharing Analyses

We used POPSTATS^37^ to calculate Outgroup *f3* statistic in the form of *f3* (Test, A, Mbuti) with Test being the Ethiopian whole-genome sequences and the NAF individuals, and A being the same set of all possible chronological and geographical proxies for the admixture. To further infer the Non-African component we used Admixtools 4.1^26^. We performed *f4* analyses using qpDstat along with the option F4:YES with this format: A,B;Test,O. As Test populations we used Ethiopian populations with non-zero contribution from the Non-African component (namely: Amhara, Somali, Wolayta and Oromo). With Admixtools we performed qpWave and qpAdm with the set of Right populations firstly defined by Lazaridis 2016, with the exception of Onge, which is not present in our analyses. Right populations used: Ust_Ishim, Kostenki14, MA1, Han, Papuan, Chukchi, Karitiana, EHG, Natufian, Switzerland_HG, WHG. We reported qpAdm results that show significance < 0.001 in qpWave, which was performed with the set of Left populations, without the Test population. We used for every analysis a custom list of Left populations to test a two-ways or a three-ways admixture. The Left populations used to perform qpAdm were selected in this order: the Test population, A and Mota for the two-ways admixture; the Test population, A, B and Mota for the three-ways admixture. Where A stands for the top scoring populations in the Outgroup *f3* analyses and B for CHG. We reported both significative and non significative results as they might be both indicative for the purpose of our analyses. We set our threshold to accept a result as significant at 0.01. We then used the information gathered from qpAdm to build a qpGraph model. We proceeded modelling qpGraph tree starting from a simple tree topology, then adding populations of interest at each step and modifying the topology to minimize the *f2* and *f4* Z-Score values.

### Bias Testing

We performed further analyses in order to detect in the unassigned sequences (X and Y components) whether important signal were lost in the deconvolution process. We compared our test populations with the *f4* statistic using this format: A,B,Test,O. As Test populations we used: Ethiopians whole genome sequences, NAF genomes, Ethiopians_J, where “J” stands for “Joint”. The Joint individuals, created for each ethnic group with Eurasian contribution (Amhara, Oromo, Somali and Wolayta), are build as a synthetic population made of the NAF and AF sequences refined by the Viterbi algorithm that passed the fbk 90% threshold, and thus not yielding the unassigned segments. To the NAF and the Ethiopians_J individuals, we added the X segments, to test if the unassigned component would give different results from the Non-African component NAF alone, which would indicate presence of biases in the deconvolution step. To the Ethiopians_J individuals along with the X component we then added the Y component as well to mimic the whole-genome. As A and B we used the possible proxy populations that may have contributed to the admixture: Levant_N, Anatolia_N, Levant_ChL. We modelled the NAF along the X component with qpAdm, using the same Left and Right populations used for the main analyses to investigate how the X component can be modelled and if the NAF with the addition of X could be modelled as the Non African component, which could indicate no bias.

