## Supplementary Information for "West Asian sources of the Eurasian component in Ethiopians: a reassessment"

### Supplementary Material

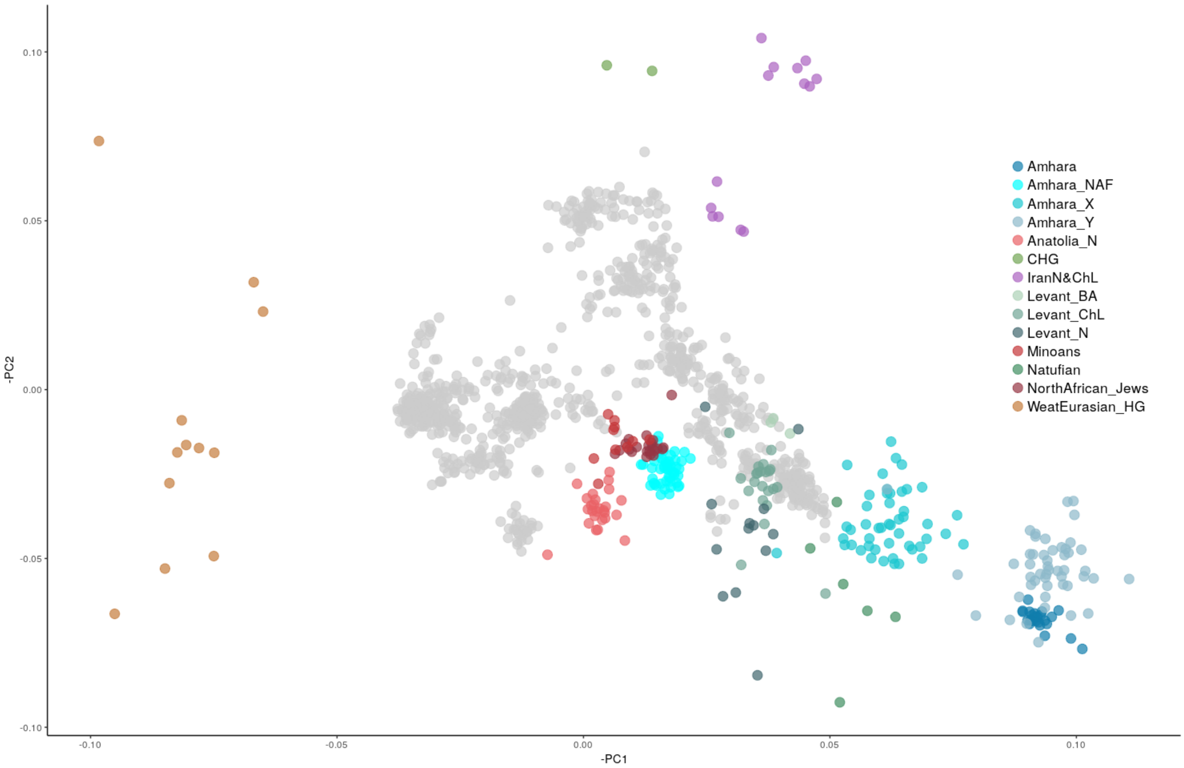


Figure S1: Principal component analysis shown in Figure 1 with the addition of the components extracted to test for biases, specifically for Amhara: X and Y (Amhara_X, Amhara_Y). The X component falls between the Near East and the Ethiopian whole-genome individuals, as expected due to the unassigned Non African sequences and the spurious African ones. Given that the Y component bears more African traces than the X component, it falls farther right in the PC1, clustering with Amhara whole-genomes. Variance explained by PC1 is 0.9% and PC2 is 0.3%

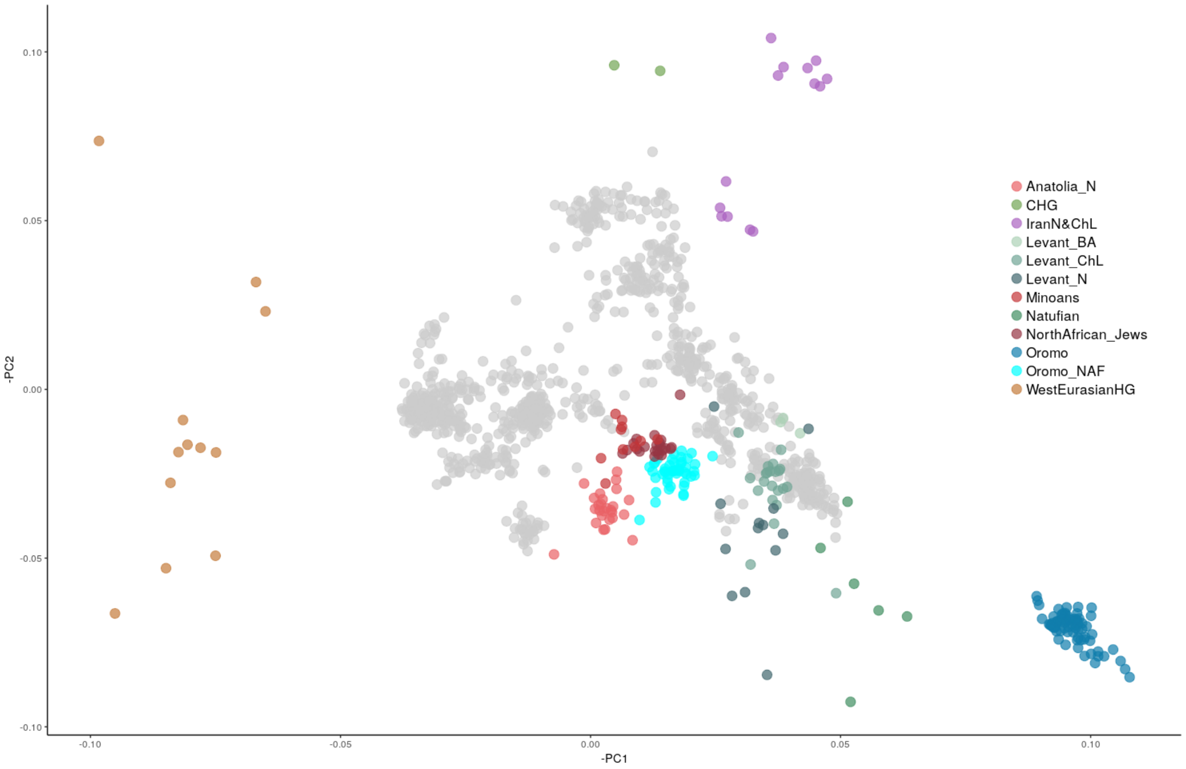


Figure S2: Principal component analysis shown in Figure 1 with Oromo population highlighted, whole-genome in dark blue and NAF sequences in light blue.Variance explained by PC1 is 0.9% and PC2 is 0.3%

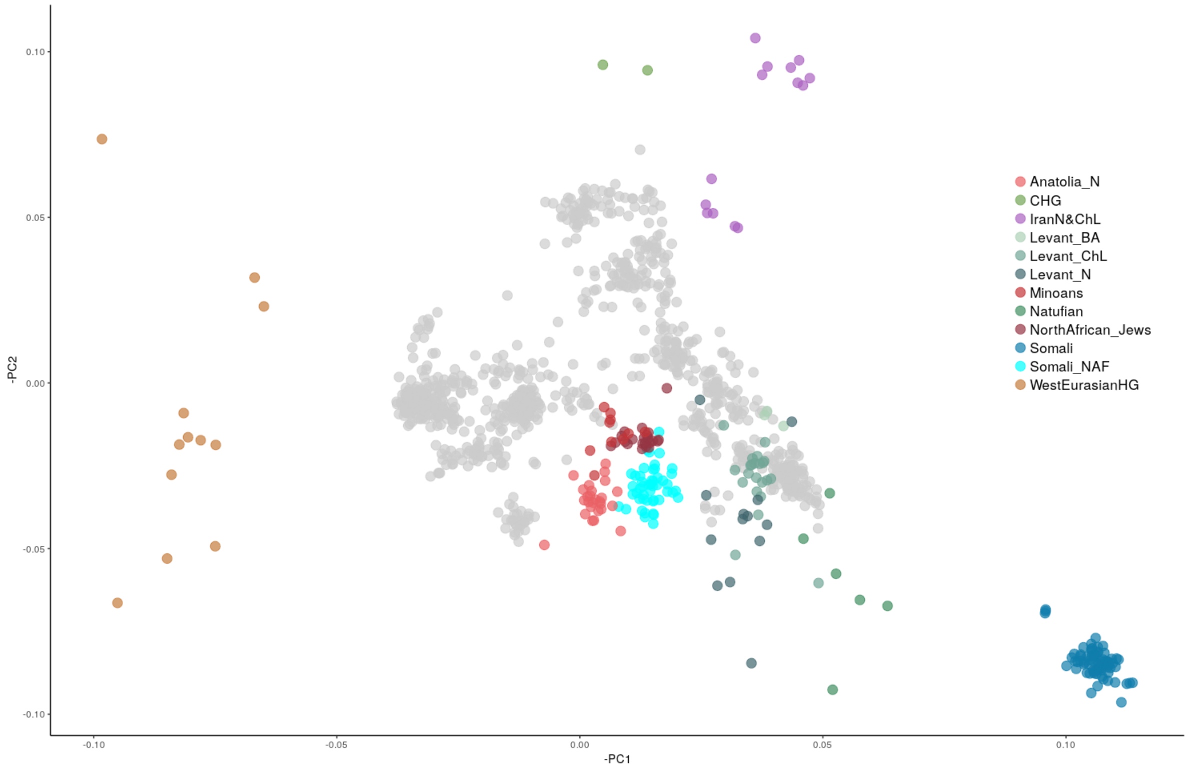


Figure S3: Principal component analysis shown in Figure 1 with Somali population highlighted, whole-genome in dark blue and NAF sequences in light blue.Variance explained by PC1 is 0.9% and PC2 is 0.3%

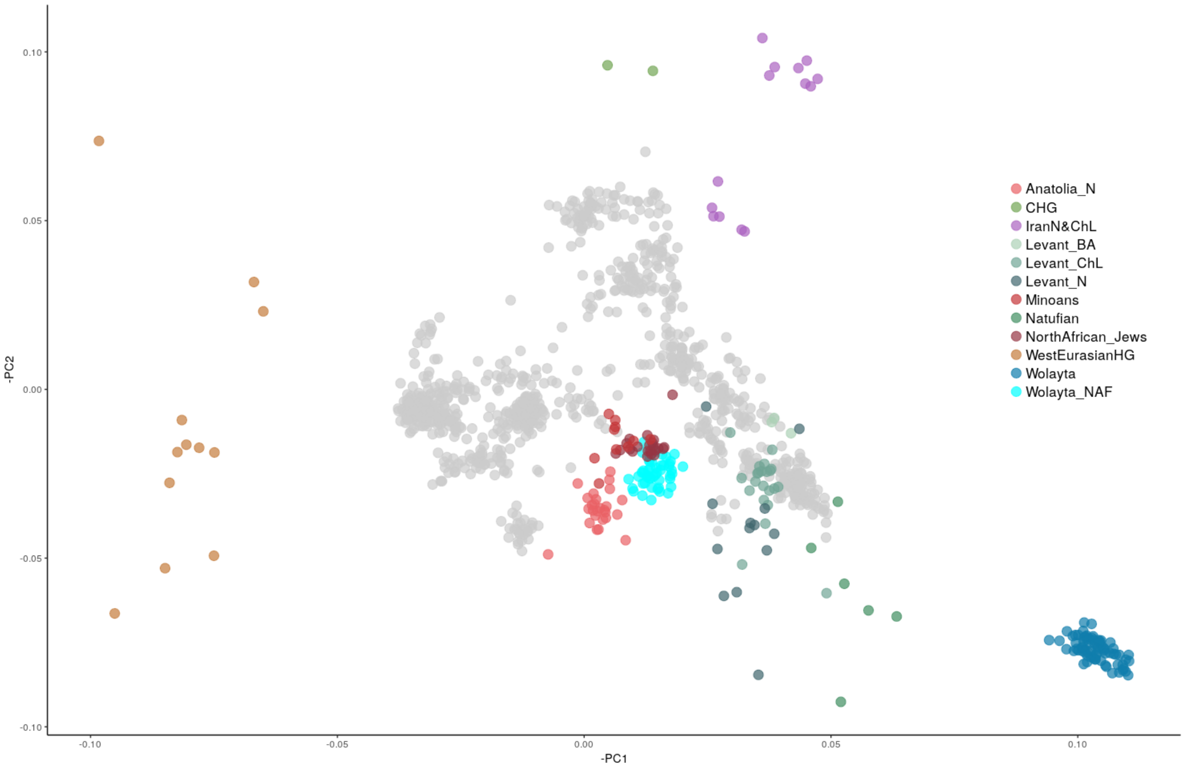


Figure S4: Principal component analysis shown in Figure 1 with Wolayta population highlighted, whole-genome in dark blue and NAF sequences in light blue.Variance explained by PC1 is 0.9% and PC2 is 0.3%

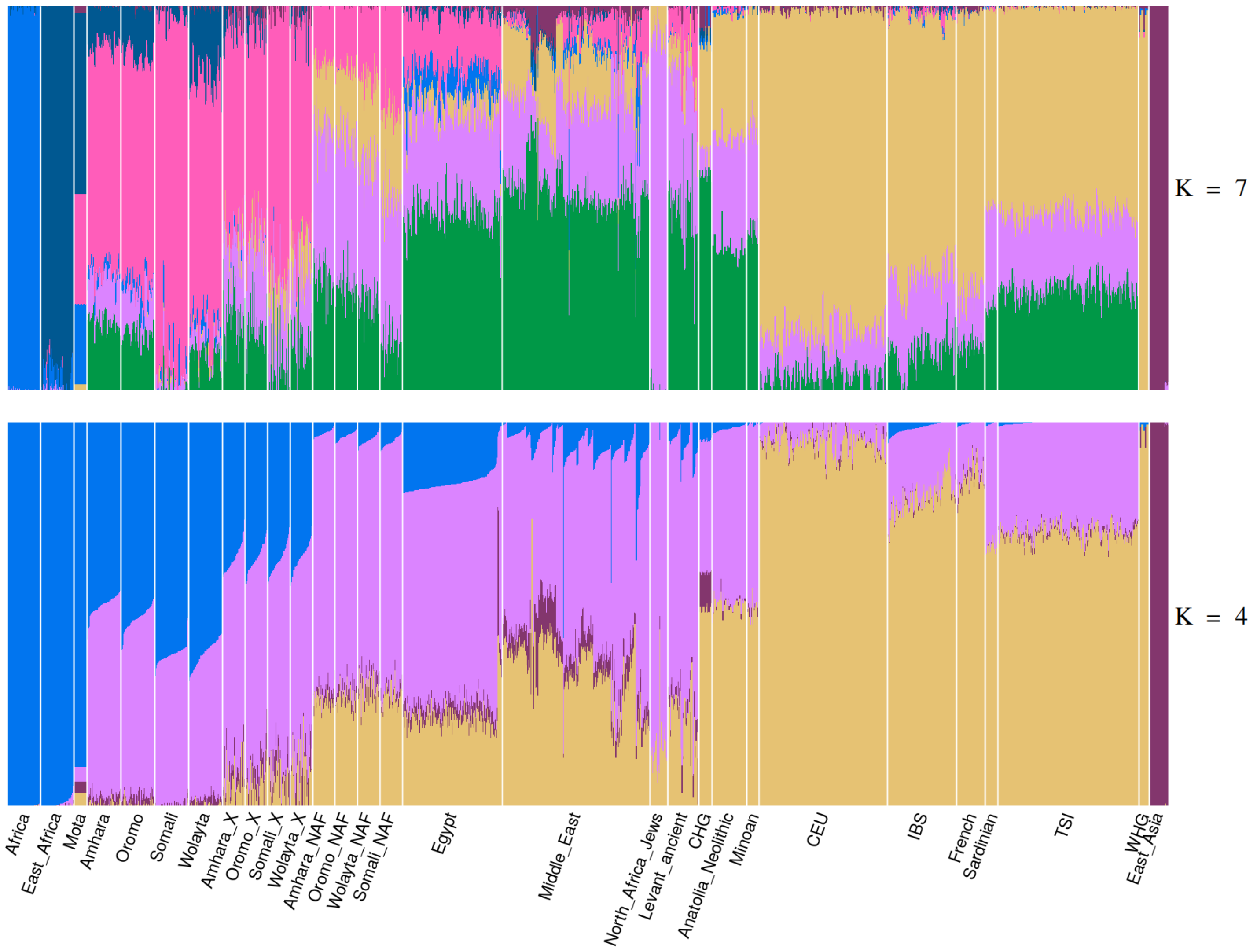


Figure S5: Supervised ADMIXTURE using modern populations as a reference on which we projected ancient and deconvoluted genomes. K=7 shows the smallest cross validation error.

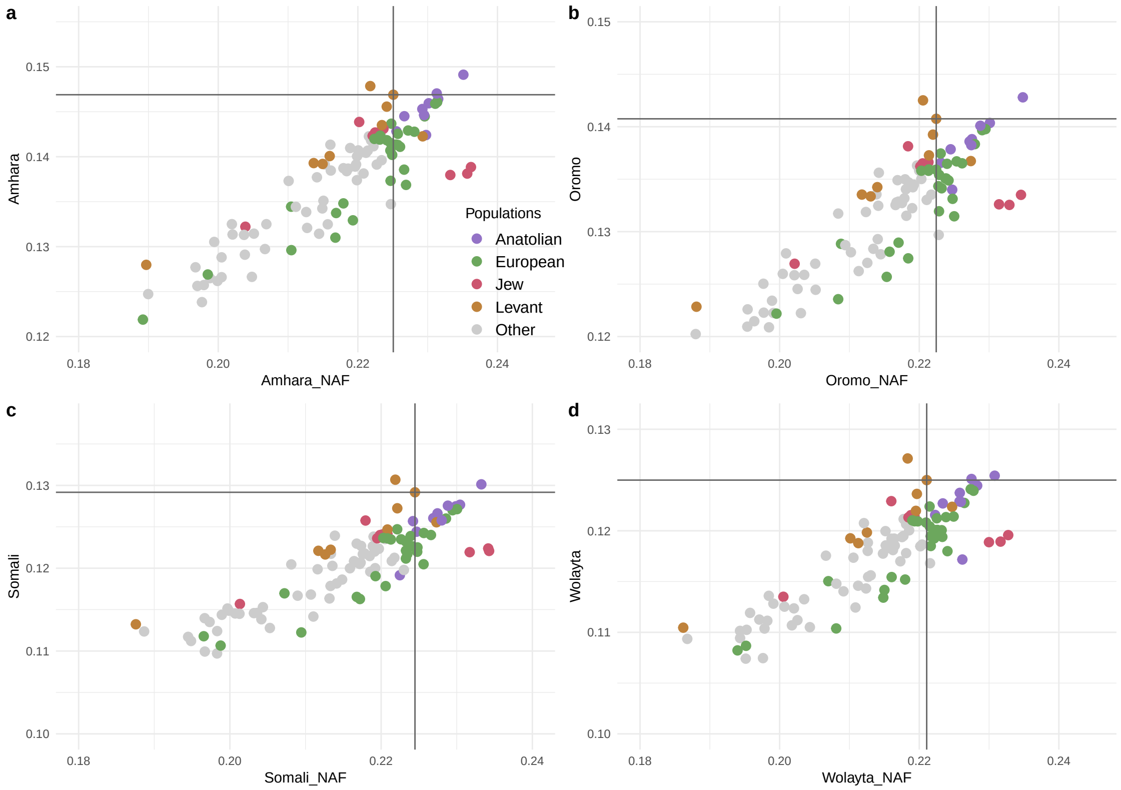


Figure S6: Scatterplot of outgroup *f3* results in form (NAF, X; Mbuti) on the x axis; *f3* in form (Amhara/Oromo/Somali/Wolayta, X; Mbuti) on the y axis, where X stands for several possible genetic and geographic neighbours. Populations are listed and colored based on their geographical origin. The vertical and horizontal lines intersect on Levant_N to highlight the most up-to-date hypothesis of the origin of the Non African component.

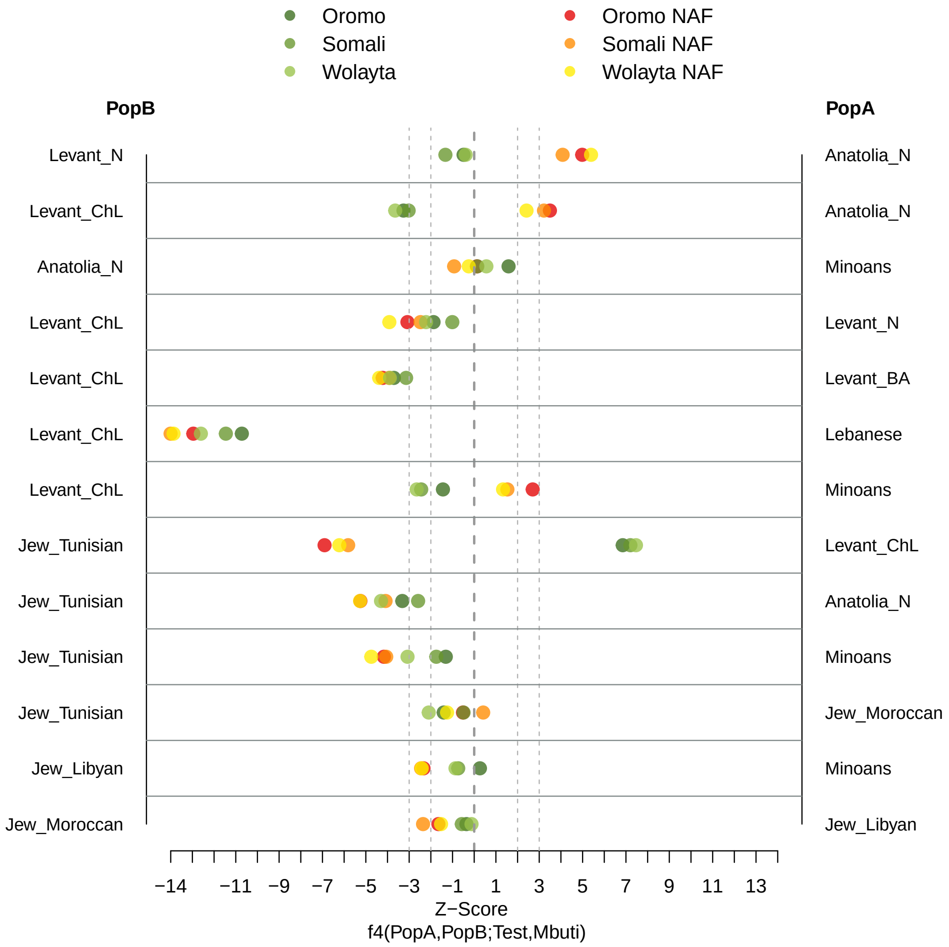


Figure S7: *f4* statistic results on Oromo, Somali and Wolayta in form of (PopA,PopB;Test,Mbuti) to test genetic similarity of the Ethiopians and respective NAF genomes to pairs of several Near Eastern populations. A and B populations are listed in the left and right side of the plot, respectively. Values in x axis indicate the Z-Scores, we draw two lines to highlight |z-Scores|= 2 and 3. Points with |z-Score| > 3 indicate a clear affinity of the test population towards one of the other population.

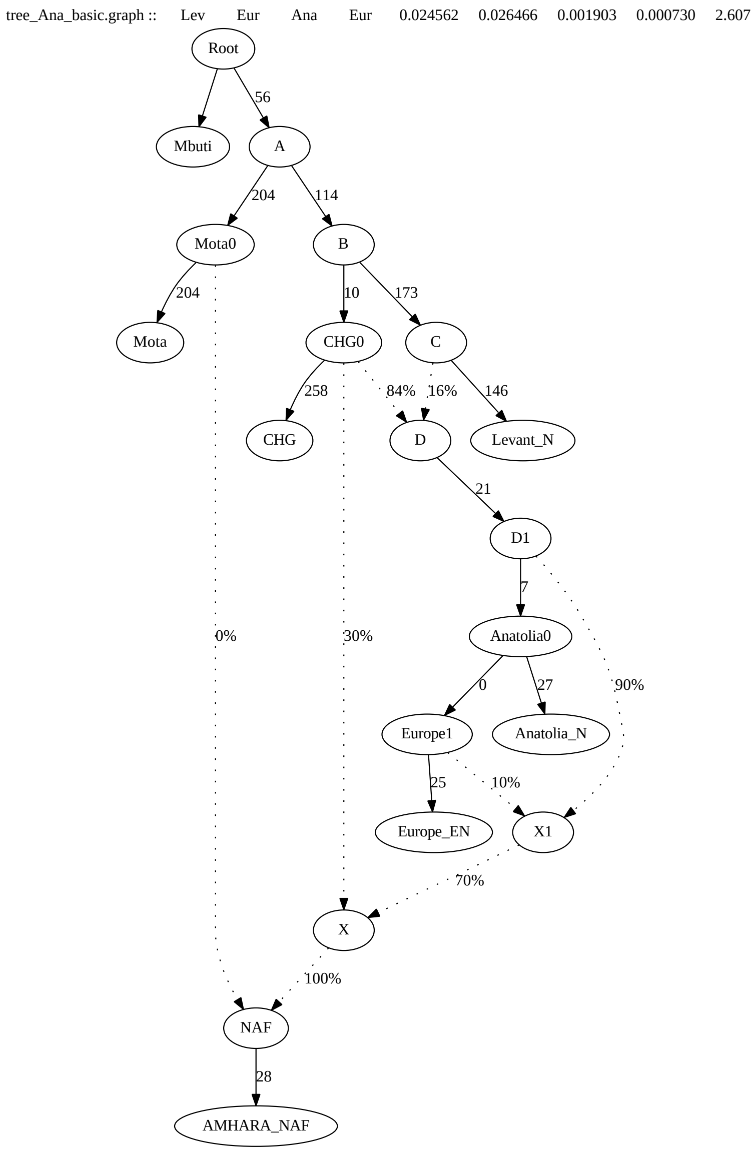


Figure S8: Admixture graph modelling Amhara NAF as being admixed with an African and two Eurasian ancestries. Worst $f4 = 2.607$, dof $= 2$ and final score $= 2149$

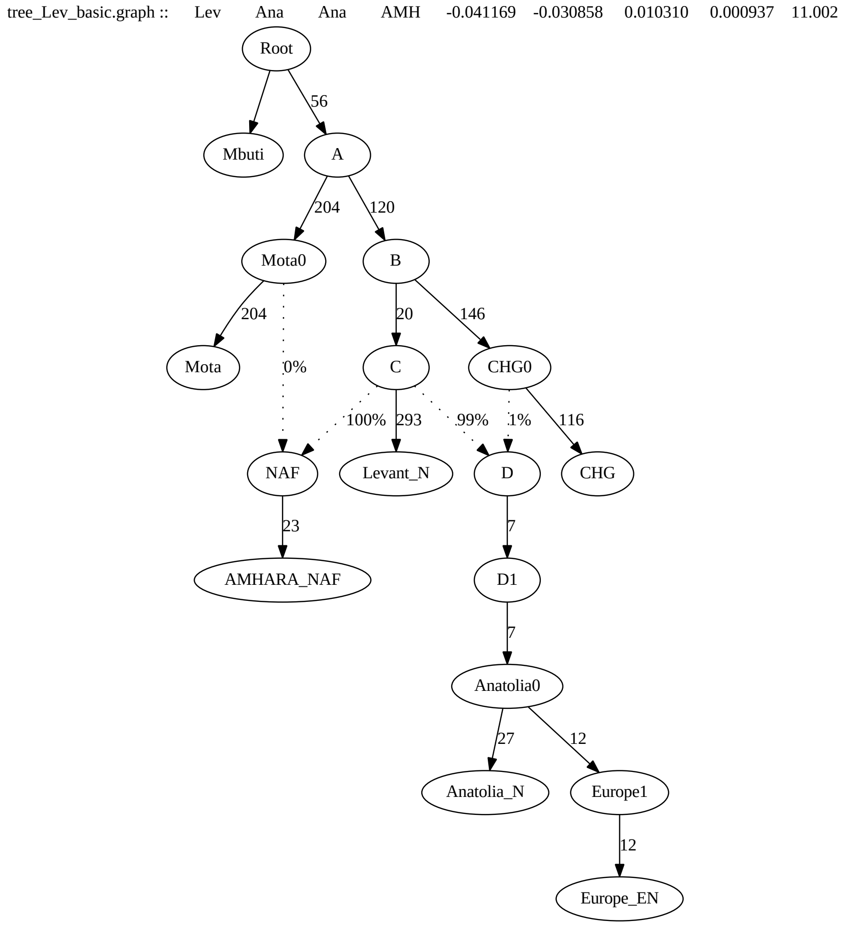


Figure S9: Admixture graph failing to model Amhara NAF as being admixed with an African and two Eurasian ancestries, one of which is Levant_N. Worst $f4$ = 11.002, dof=4 and final score = 16522

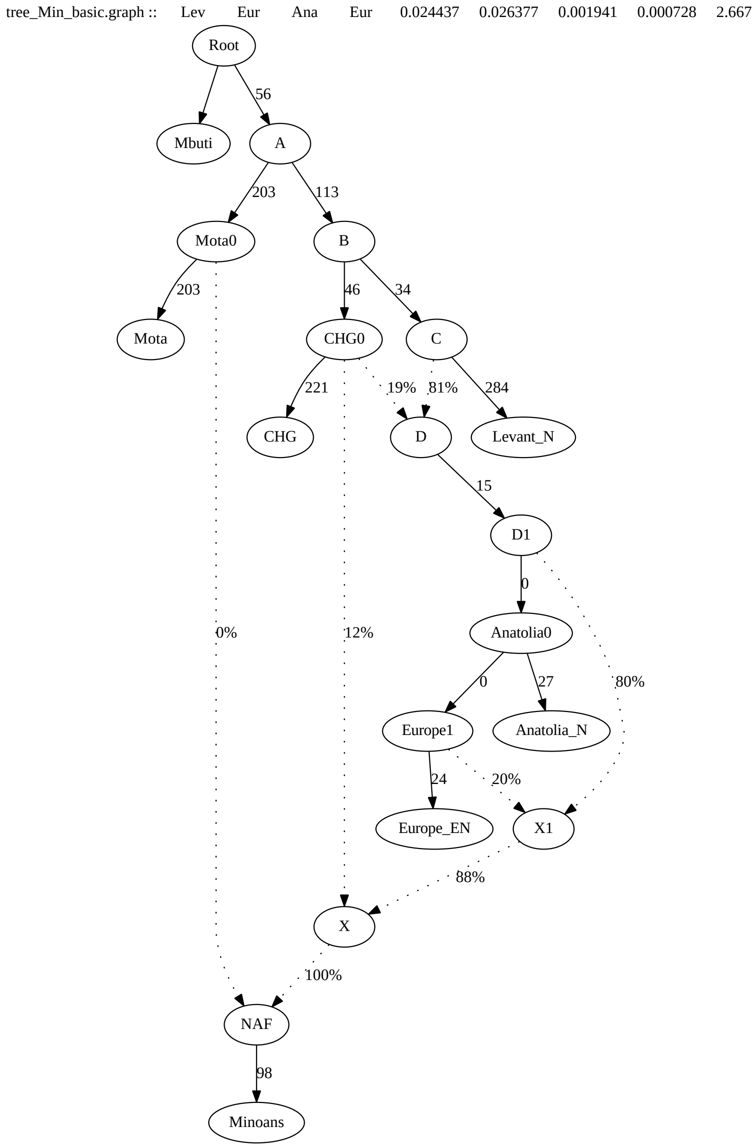


Figure S10: Admixture graph modelling Minoans as being admixed with an African and two Eurasian ancestries. Worst $f4$ = 2.667, dof = 2 and final score = 2046.

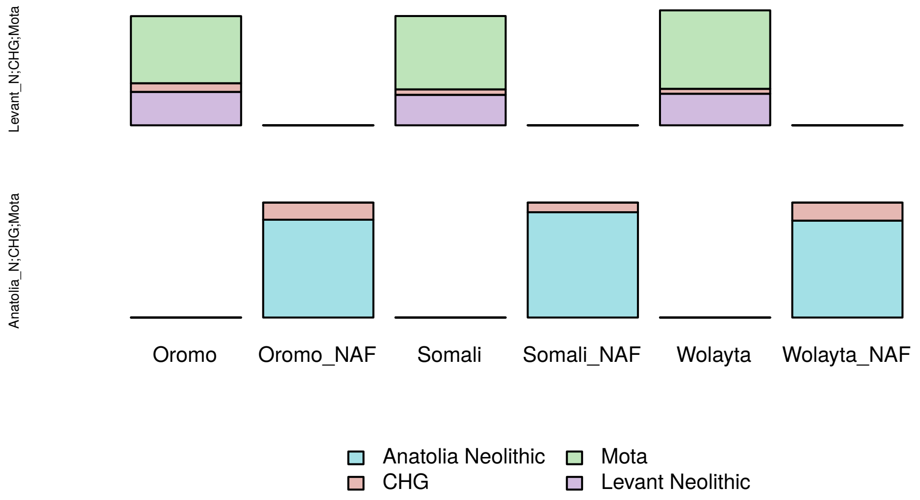


Figure S11: Modelling Oromo, Somali, Wolayta and their respective NAF component as a mix of Mota and Near Eastern populations, with 2 and 3 ways admixtures. Violet indicates the Levant_N component, pink the Central Hunter-Gatherers, light green the African component and light blue highlights the Anatolian ancestry. The left side of the graph lists the sources used to model the populations in the x axis; unfilled boxes indicate unfeasible results or p-value $< 0.01$.

Table S1: F4 results in form (PopA,PopB; Test,O) testing genetic similarity between Ethiopians (Test) and different Near Eastern populations (PopA and PopB). See Supplementary Table S1 provided as a separate file.

Table S2: Modelling Ethiopians and North African Jews as a mixture of Mota and different Near Eastern populations. P-values $> 0.05$ are indicated in bold. See Supplementary Table S2 provided as a separate file.
